## Supplementary Figure 1 for "Intracellular accumulation of free cholesterol in macrophages triggers a PARP1 response to DNA damage and PARP1 impairs lipopolysaccharide-induced inflammatory response"

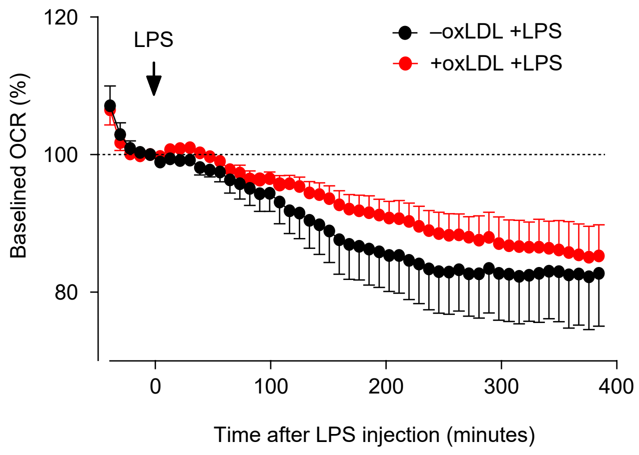


**Characterizing the effect of oxLDL loadeding on the metabolic profile of PMφs after LPS stimulation.**

Real-time OCR measurements were performed using a Seahorse analyzer and PMφs with or without oxLDL accumulation (±oxLDL) and LPS stimulation. OCR measurements were normalized to one reading cycle prior to LPS injection (indicated by an arrow). The mean ± SEM is plotted (n = 4). Significant differences between corresponding time points in –oxLDL versus +oxLDL groups were not detected using a one-way ANOVA and a Bonferroni post hoc test.
