## Supplementary Figure 2 for "Intracellular accumulation of free cholesterol in macrophages triggers a PARP1 response to DNA damage and PARP1 impairs lipopolysaccharide-induced inflammatory response"

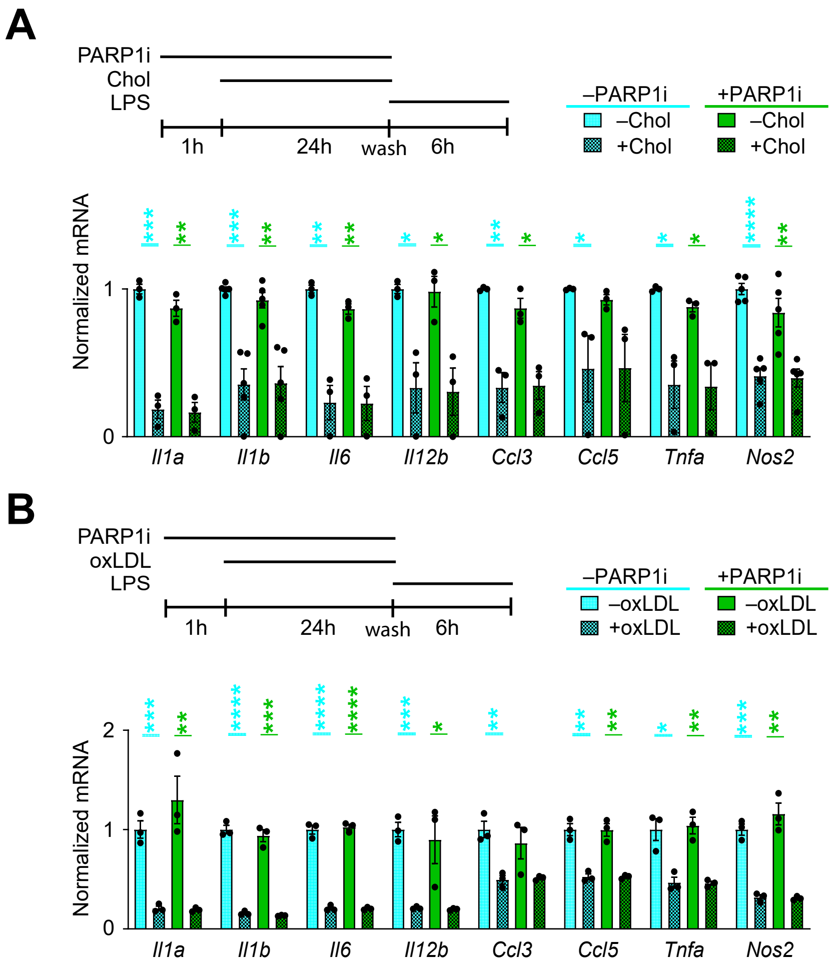


**Inhibition of PARP1 enzymatic activity does not rescue the suppression of LPS-induced inflammatory gene expression in lipid loaded PMφs.**

**(A, B)** AG14361, a PARP1 inhibitor (PARP1i), was added 1 h prior to culturing PMφs with or without free cholesterol (Chol, A, n = 3-5) or oxLDL (B, n = 3-5). Subsequently PMφs were stimulated with LPS for 6 h, as is shown in a schematic above each graph. Inflammatory gene expression was measured by qPCR. The mean ± SEM is plotted. Significant differences were determined using an unpaired Student's t test (* $P < 0.05$, ** $P < 0.01$, *** $P < 0.001$, **** $P < 0.0001$).
